# Illusory cold desensitizes touch

**DOI:** 10.64898/2026.06.02.729707

**Authors:** Juan Carlos Ramirez, Jose Vergara, Brandon Kim, Hannes P. Saal, Jeffrey M. Yau

## Abstract

In Jack London’s *To Build a Fire*, the doomed protagonist notes that “the tremendous cold had already driven the life out of his fingers… dead fingers could neither touch nor clutch”. Indeed, our hands are numbed when exposed to cold, a desensitization process traditionally attributed to biophysical and physiological effects on mechanoreceptor and afferent responses^1^. Yet, perception is not based solely on sensory inputs and often reflects prior beliefs and expectations^2^. Accordingly, the mere experience of coldness, given its association with tactile desensitization, conceivably modulates touch independent of physical effects. We dissociated the physical and subjective influences of cooling on touch by leveraging the Thermal Referral Illusion^3–7^, in which cooling of the index and ring fingers induces an illusory perception of coldness that is referred to the middle finger which did not experience local cooling. We found that both veridical and illusory cold elevated tactile detection thresholds. Threshold shifts scaled with subjective ratings of cold intensity which varied with thermal adaptation. Thermal referral induction did not result in changes to skin temperature, ruling out indirect physical effects on the tested finger. To account for cold effects on touch, we implemented a computational model incorporating both physical and subjective factors by simulating changes in tactile afferent spiking activity^8^ and modulation of decision computations, respectively. The model reproduced empirical cooling effects on afferent responses^9^ and recapitulated psychophysical detection patterns. Our results reveal that subjective thermal perception contributes to cold-induced attenuation of touch, highlighting how multimodal experiences and expectations shape somatosensation.

## Results

### Veridical and illusory cold are similarly robust and durable

Thermal perception adapts rapidly in response to sustained exposure to a temperature state^10,11^. Accordingly, we first quantified the magnitude and durability of veridical and illusory thermal percepts induced by our thermal stimulation patterns (Figure 1A) before characterizing thermotactile interactions. Independent thermoelectric Peltier devices^12^ contacted the distal pads of the index, middle, and ring finger on each participant’s left hand to deliver well-controlled and stable thermal cues over 10-min adaptation periods (Figure S1). For baseline (neutral) tests, all three fingers were maintained at 30°C. Under the veridical cold condition, all three fingers experienced 20°C stimulation. Under the illusory condition, the index and ring fingers experienced 20°C while the flanked middle finger was maintained at 30°C. Participants reported the perceived thermal intensity on their middle finger every 20 seconds (Methods), producing 30 ratings for each thermal condition (neutral, veridical, illusory) over the adaptation periods. Following the adaptation periods, all three fingers immediately experienced 30°C for an additional 10-min period. Beyond simply readapting participants to thermal neutral, testing during these periods enabled us to quantify the strength and duration of aftereffects induced during the adaptation periods.

**Figure 1.**
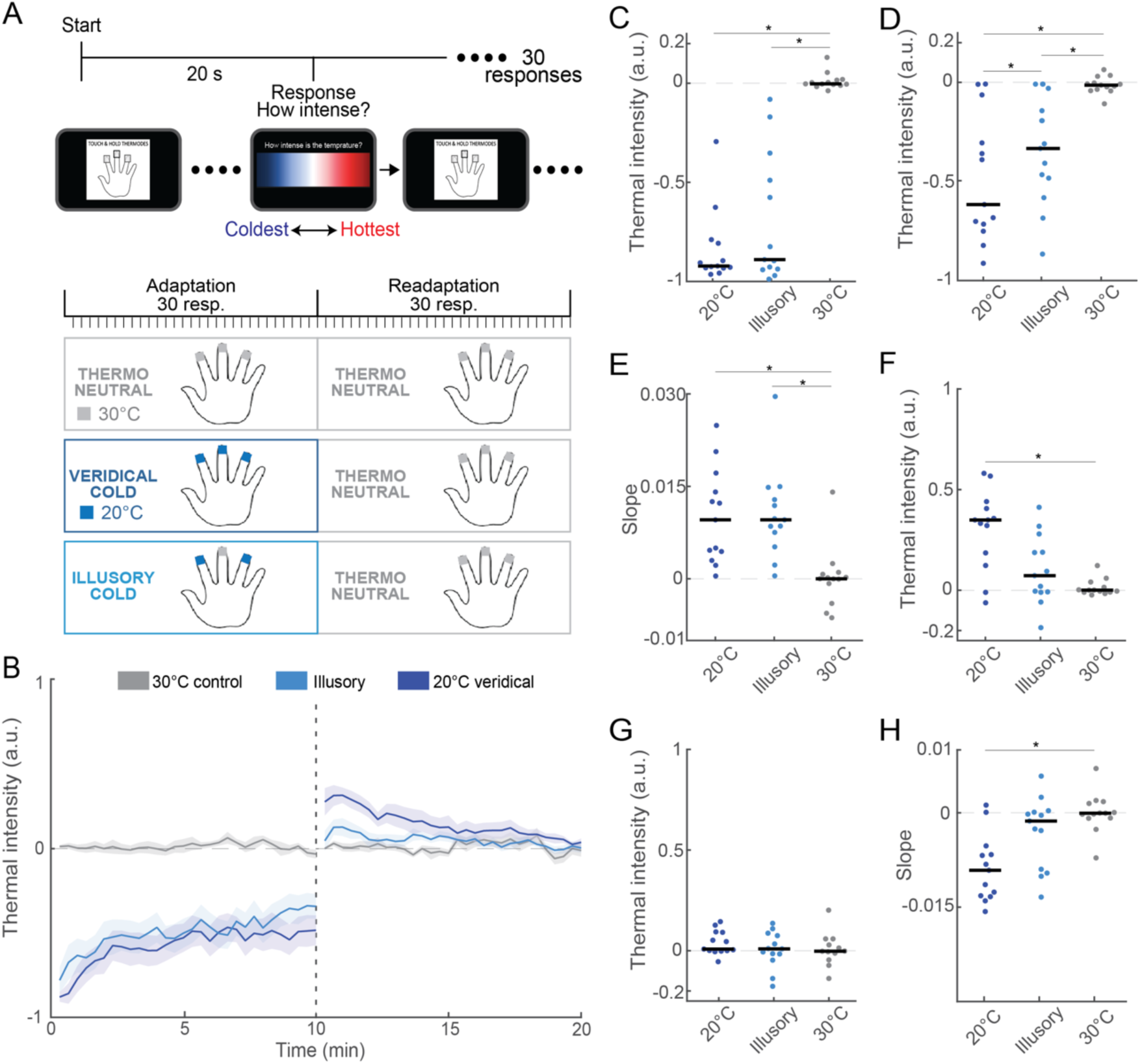
Illusory and veridical cold percepts are similarly intense and durable. (A) Thermal adaptation task design. Participants experienced thermal cues on their index, middle, and ring fingers. Participants reported the perceived thermal intensity on their middle finger every 20 seconds over 10-min adaptation periods during which they experienced neutral stimulation (30°C on all fingers), veridical cold (20°C on all fingers), and illusory cold (30°C on middle finger, 20°C on flanking fingers). Immediately following the adaptation periods, participants reported the perceived thermal intensity on their middle finger while experiencing a 10-min readaptation period comprising 30°C stimulation on all fingers. (B) Traces depict normalized thermal intensity ratings (mean ± SEM, n = 13) across adaptation and readaptation periods under each thermal condition. Positive/negative intensity ratings correspond to hot/cold percepts. (C) Mean thermal intensity ratings during the first minute of adaptation. Markers indicate individual participant ratings. (D) Mean thermal intensity ratings during the final minute of adaptation. Conventions as in C. (E) Slope values quantifying rate of change of intensity values over adaptation period. Positive slopes indicate reduced cold intensity ratings over time. Black lines indicate group averaged slope. Markers indicate individual participant slopes. (F) Mean thermal intensity ratings during the first minute of readaptation. Conventions as in C. (G) Mean thermal intensity ratings during the final minute of readaptation. Conventions as in C. (H) Slope values quantifying rate of change of intensity values over readaptation period. Negative slopes indicate reduced heat intensity ratings over time. Conventions as in E.

Thermal intensity ratings revealed robust cold percepts that evolved gradually over the veridical and illusory cold adaptation periods (Figure 1B). Thermal intensity ratings (negative values indicate cooler) during the first minute of adaptation (Figure 1C) differed significantly across conditions (Friedman test; χ²(2) = 19.54, p = 5.72×10^-5^). Veridical (−0.84±0.05) and illusory (−0.70±0.01) cold produced similar thermal ratings (signed-rank test; mean rank difference = 0.08, 95% CI [–0.86, 1.02], p = 1, W = 36) that each differed significantly from neutral condition ratings (0.01±0.01) (signed-rank test; veridical vs neutral: mean rank difference = –1.46, 95% CI [–2.40, –0.52], p = 5.83×10^-4^, W = 0; illusory vs neutral: mean rank difference = 1.54, 95% CI [0.60, 2.48], p = 2.63×10^-4^, W = 0). Significant differences in thermal ratings persisted into the final minute of the adaptation period (Figure 1D; rmANOVA; F(2, 24) = 20.64, p = 6.10×10^-6^), even as perceived intensity generally declined in both cold conditions. At the end of the adaptation period, participants continued to judge both the veridical (t = 5.12, p = 7.54×10^-4^) and illusory (t = 4.19, p = 3.75×10^-3^) conditions as more intense compared to the neutral condition (−0.01±0.01). Veridical cold (−0.49±0.09) also tended to be judged as more intense (95% CI [−0.68, −0.27], t = 2.90, p = 0.04) compared to illusory cold (−0.35±0.07). The reduced thermal intensity ratings in the final minute are consistent with adaptation^11^, which we quantified by fitting linear functions (Methods). Adaptation rates (Figure 1E) differed significantly across conditions (Friedman test; χ²(2) = 14.92, p = 5.75×10^-4^). Adaptation with veridical cold (0.01±0.002; mean rank difference = 1.08, 95% CI [0.14, 2.02], p = 0.02, W = 89) and illusory cold (0.01±0.002; mean rank difference = −1.46, 95% CI [−2.40, −0.52], p = 5.83×10^-4^, W = 90) was greater than that observed in the neutral condition (1.6×10^-4^ ±0.0014). Adaptation rates did not differ between the cold conditions (mean rank difference = −0.38, 95% CI [−1.32, 0.55], p = 0.98, W = 32). Collectively, these results indicate that participants experienced the illusory cold condition comparably to the veridical cold condition. Furthermore, the adaptation time courses for the veridical and illusory cold percepts validate that thermotactile interactions can be reliably assessed within a 10-min period.

Sustained exposure to sensory inputs leads to adaptation in peripheral and central neurons^13,14^. Such adaptation can produce perceptual aftereffects in the opposite direction when downstream decoding mechanisms do not also adapt^15–17^. To characterize the strength and duration of thermal aftereffects, participants rated the intensity of the thermoneutral 30°C stimulation pattern over a 10-min readaptation period following adaptation to the cold and neutral conditions (Figure 1A). Given the adaptation observed with veridical and illusory cold, we predicted that participants would subsequently rate the neutral condition as feeling warm. Consistent with this prediction, thermal ratings tended to be positive immediately following the switch to uniform 30°C stimulation (Figure 1B). Intensity ratings during the first minute of readaptation differed significantly according to the adaptation conditions (Figure 1F; Friedman test; χ²(2) = 8.0, p = 0.02). While positive intensity ratings followed adaptation to both veridical (0.30±0.05) and illusory cold (0.10±0.05), only the ratings following veridical cold adaptation differed significantly (mean rank difference = −1.08, 95% CI [−2.01, −0.14], p = 0.02, W = 3) from ratings following the neutral condition adaptation (0.01±0.01). By the end of the readaptation period (Figure 1G), thermal intensity ratings were indistinguishable across conditions (Friedman test; χ²(2) = 2.0, p = 0.37), indicating that participants had readapted fully. We quantified readaptation rates under each condition (Figure 1H) and found a significant effect of adaptation condition (Friedman test, χ²(2) = 12.92, p = 1.56×10^-3^), driven by differences (mean rank difference = 1.38, 95% CI [0.45, 2.32], p = 1.25×10^-3^, W = 90) in readaptation rates between the veridical cold (−8.85×10^-3^±0.002) and neutral conditions (1.73×10^-5^±8×10^-4^). Readaptation rates following illusory cold adaptation (−3.12×10^-3^±0.002) did not differ from readaptation rates in any other condition. These results reveal the context-dependent nature of thermal aftereffects, which rely more on adaptation driven by physical temperature changes rather than the subjective experience of cold^7^. Critically, complete washout occurs with 10-min exposure to 30°C stimulation indicating that thermotactile interactions can be fairly assessed assuming appropriate readaptation periods.

### Veridical and illusory cold both elevate vibration detection thresholds

Veridical cold has been shown to elevate tactile detection thresholds^3,9, 18–25.^ Whether this thermotactile interaction results solely from physical changes in the skin or reflects modulation by the subjective experience of cold has not been determined. Having established the magnitude and durability of veridical and illusory cold percepts, we characterized the effect of veridical and illusory cold on tactile sensitivity, dissociating the physical and subjective thermal influences on touch. To quantify each participant’s tactile sensitivity, we determined their vibration detection thresholds using a 2-interval, 2-alternative forced-choice paradigm (Figure 2A). On each trial, participants received a 250-Hz vibration on their middle finger during 1 of 2 trial intervals. Participants were tasked with reporting the trial interval in which they detected the vibration cue. Vibration amplitude was determined using an adaptive staircase procedure (Methods): If the participant correctly identified the interval containing the cue on 2 trials, vibration amplitude on the subsequent trial was reduced. Conversely, if the participant reported detecting the cue in the incorrect interval on a single trial, vibration amplitude on the subsequent trial was increased. A staircase terminated after 10 amplitude reversals (28.92±0.33 total trials) and the average amplitude over the final 5 trials was computed for a threshold estimate. Three staircases (2 descending, 1 ascending) were completed serially under each condition. Participants were tested in a full factorial design (Figure 2B) comprising 3 thermal states (neutral, veridical cold, and illusory cold) and 2 vibration durations (100 and 500ms) resulting in 6 total conditions (18 staircases) per participant. The inclusion of the duration manipulation served to test whether thermal modulation of touch operated independently or interactively with another stimulus feature known to influence vibration detection thresholds ^26–29^. To ensure participants experienced each thermal condition from a thermal neutral state, the index, middle, and ring fingers were readapted to 30°C for 10 minutes prior to threshold testing.

**Figure 2.**
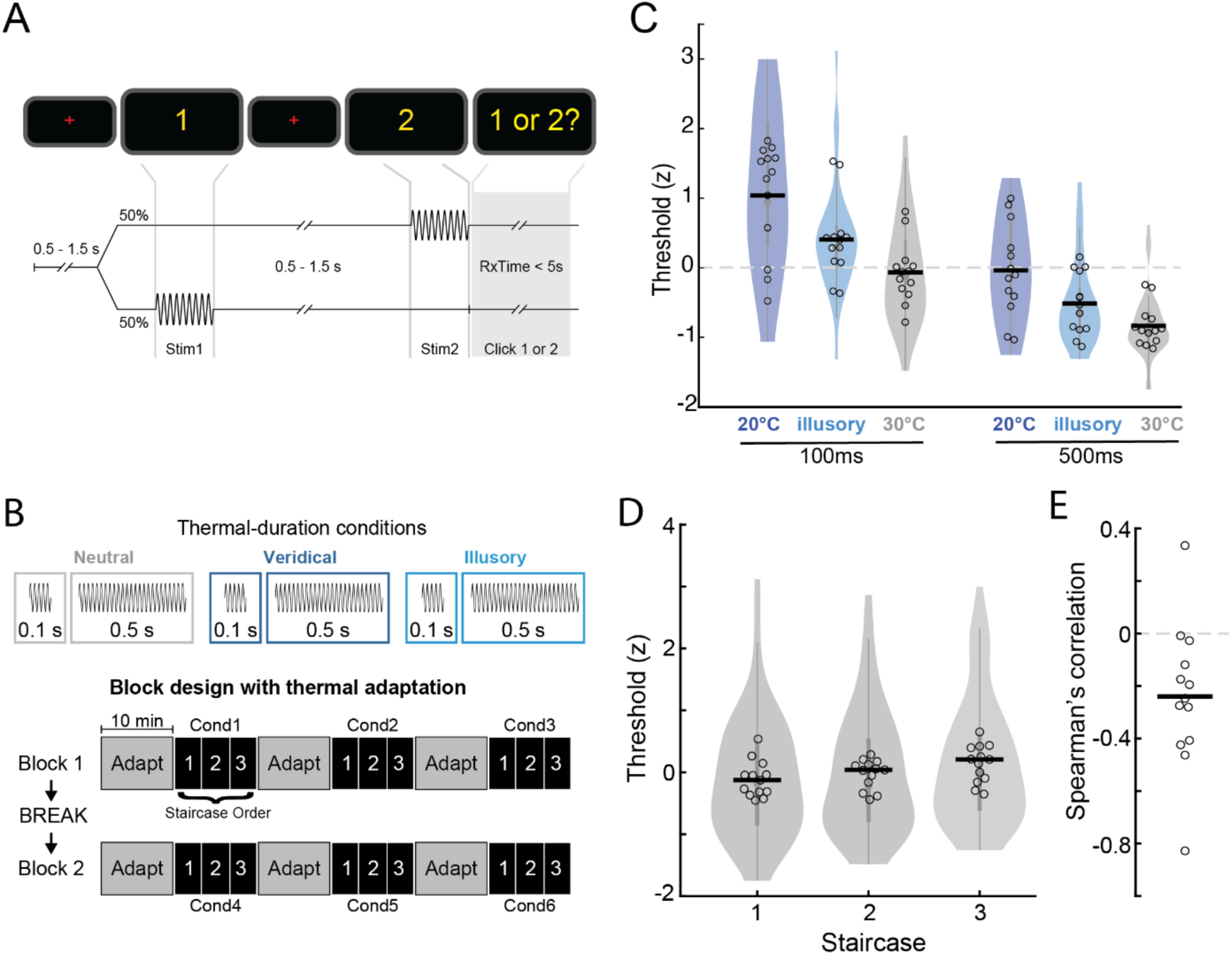
Illusory and veridical cold elevate tactile detection thresholds. (A) Tactile detection task. A peri-threshold tactile stimulus (250-Hz vibration) is delivered to the middle finger in one of two stimulus intervals on each trial. Participants are tasked with reporting whether they detected the stimulus in the first or second interval. Stimulus amplitude is determined according to a 2-down/1-up adaptative staircase procedure. Stimulus amplitudes in the ascending and descending staircases start at 0.01mm and 3mm, respectively. (B) Tactile detection thresholds are tested under 6 experimental conditions comprising 3 temperature conditions (neutral, veridical cold, illusory cold) and 2 vibration durations (100 and 500ms). Three staircases (2 descending and 1 ascending) are performed for each condition after a 10-min adaptation period to 30°C stimulation on the index, middle, and ring fingers. (C) Normalized vibration detection thresholds under 6 thermal and duration conditions. Horizontal lines indicate group medians. Circles indicate individual participant’s thresholds averaged over staircases. Violin plots show threshold distributions estimated from individual staircases. (D) Vibration detection thresholds in each staircase marginalized over temperature and duration conditions. Conventions as in C. (E) Relationship between vibration detection thresholds and thermal intensity ratings. Negative Spearman’s correlation values indicate higher thresholds are associated with more intense cold ratings. Horizontal line indicates group average. Circles indicate Spearman’s correlations computed for individual participants.

Figure 2C shows detection thresholds sorted by thermal and duration conditions. These threshold patterns revealed significant main effects of cue duration (F(1, 204) = 94.43, p = 2.22×10^-16^) and thermal condition (F(2, 204) = 32.23, p = 6.89×10^-13^). No other main or interaction effects (Methods) achieved statistical significance (all *p*s > 0.05). As shown previously^26–29^, detection thresholds were higher with shorter vibration cues. Across durations, thresholds were significantly higher (t = 8.03, p < 1×10^-4^) with veridical cold compared to thresholds measured under the thermoneutral condition, consistent with prior reports^30^. Notably, thresholds measured under illusory cold were also significantly elevated compared to thermoneutral thresholds (t = 4.00, p = 3×10^-4^). This was a reliable pattern observed in either or both duration conditions in 12 of 13 participants, significantly greater than chance levels (binomial test, p = 1.71×10^-3^). The significant influence of illusory cold demonstrates that direct cooling of the skin is not required for cold-mediated tactile sensitivity changes. Veridical cold induced larger (t = 4.03, p = 2×10^-4^) threshold changes (3.9 dB relative to thermoneutral) compared to illusory cold (2.7 dB relative to thermoneutral) implying that physical and subjective factors distinctly contribute to thermotactile interactions.

Although thresholds did not differ significantly between the different staircase procedures (Figure 2D; F(2,204) = 2.88, p = 0.06), we considered whether threshold variations related to the *perceived* thermal state of the middle finger, which adapted gradually with 10-min sustained exposure to the cold conditions. To relate detection thresholds to thermal intensity ratings, we computed each participant’s mean intensity rating over 1-min intervals beginning in the 3^rd^, 6^th^, and 9^th^ minute of the adaptation periods (Methods). These intervals corresponded to the average times when participants completed the final 5 trials of the staircase which were used for computing detection thresholds under each condition. We found that thresholds were negatively correlated (r = −0.24±0.08) with thermal intensity ratings and differed significantly from 0 (Figure 2E; t = −3.13, p = 8.64×10^-3^). Higher intensity ratings corresponded to higher detection thresholds in 12 of 13 participants, significantly greater than chance levels (binomial test, p = 1.71×10^-3^). Given that the physical temperature of the thermoelectric coolers and test finger (Figure S2) remained stable over the test periods, these results further highlight the role of subjective thermal perception in thermotactile interactions.

### Tactile attenuation with illusory cold is not due to vasoconstriction or cardiac phase effects

By what mechanism might the subjective experience of cold modulate tactile sensitivity? We considered the possibility that cooling of the index and ring fingers – while maintaining the thermoelectric contact on the middle finger at 30°C – may have triggered a physiological reaction that altered the transduction and signaling of vibrations on the middle finger during the illusory cold condition. Indeed, cooling of the body can induce vasoconstriction in the extremities^31^. Extreme cooling of one hand can even induce vasoconstriction in the other hand as measured by changes in skin temperature^32^. These indirect, reflexive responses, mediated by sympathetic adrenergic nerves, are considered to be thermoregulatory processes that reduce heat loss and maintain core temperature in a metabolically efficient manner^33^. In separate experiments, we tested for indirect vasoconstrictive effects on the middle finger by measuring the temperature of the unloaded middle finger when the flanking fingers received either 20°C or 30°C thermal stimulation (Methods). If the illusory condition was associated with vasoconstriction of the middle finger, we predicted that the middle finger’s temperature would be lower when the flanking fingers were maintained at 20°C. Inconsistent with this prediction, we found the unloaded middle finger’s temperature did not change significantly (F(2,10) = 6.7×10^-3^, p = 0.94) as the flanking fingers transitioned from thermoneutral to cold and back to thermoneutral (Figure S3). These results argue against elevated middle-finger thresholds under illusory cold being caused by indirect vasoconstriction.

The ability to detect a faint tactile cue can depend on the phase of the cardiac cycle in which the stimulus falls^34,35^, with better detection for stimuli falling within the systolic phase. In fact, active tactile sampling is adjusted with respect to the cardiac cycle to maximize sensitivity^36^. Even though our experimental design prevented participants from anticipating vibration presentations to strategically time their evidence accumulation, we evaluated the effects of thermal and duration manipulations while accounting for the influence of cardiac cycle. Specifically, we determined whether the amplitudes of the peri-threshold stimuli over all tested staircases differed according to cardiac cycle phase (Figure S4) in addition to temperature and duration (Methods). Consistent with their influence on threshold estimates, we found significant main effects of thermal condition (F = 7.71, p = 4.83×10^-4^) and duration (F = 9.19, p = 2.52×10^-3^) on stimulus amplitudes. Notably, the main effect of cardiac phase and all interaction effects failed to achieve statistical significance (all ps > 0.05). These results indicate that the thermotactile effects on vibration detection thresholds are independent of cardiac-related sensory attenuation.

### A computational detection model based on simulated Pacinian afferent activity reproduces cold attenuation effects

Our psychophysical results show that detection thresholds are elevated when the finger is experienced as cold due to both physical and subjective factors. We next considered the neural basis of the physical factors underlying thermotactile interactions. Although the skin contains multiple classes of mechanoreceptors with overlapping response properties^37^, Pacinian corpuscles are the only credible candidates in our experiment due to their exceptional sensitivity to high-frequency vibrations^38,39^, especially at low amplitudes, with all other receptor thresholds at least an order of magnitude higher at our test frequency of 250Hz^40,41^. Additionally, the response profiles of Pacinians have been shown to depend on thermal conditions^9,42,43^. To test the role of Pacinian afferents in temperature-modulated vibration detection, we used a biologically plausible skin-neuron model of the hand, TouchSim^8^, to simulate spiking activity in Pacinian afferent populations and model vibration detection behavior (Figure 3A).

**Figure 3.**
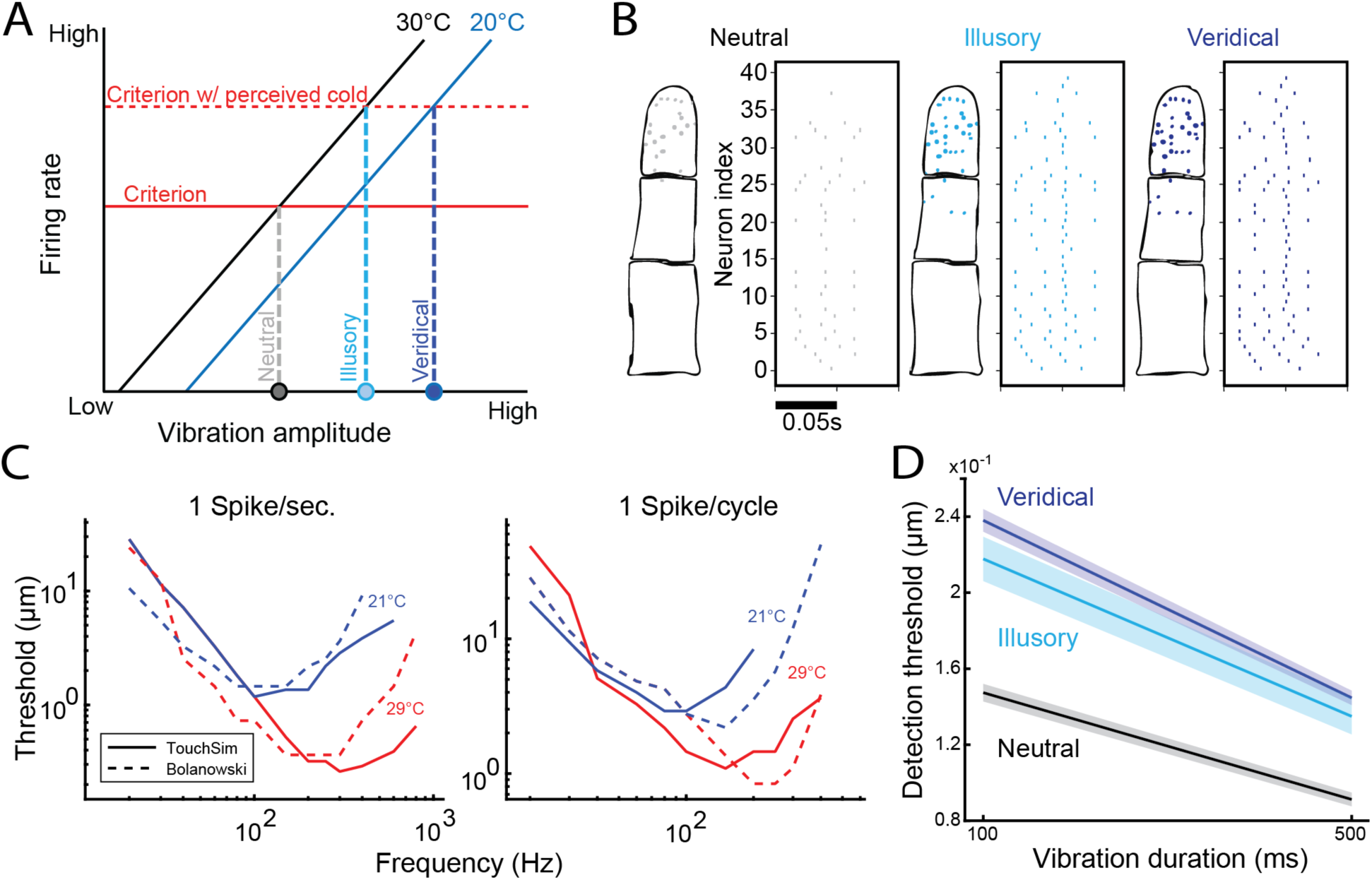
Thermotactile interactions result from cold-modulation of afferent spiking rates and decoding rule. (A) Two-factor model of thermotactile interactions. Thermoneutral response function (30°C; black line) indicates that increases in vibration amplitude are associated with increasing spiking activity in the afferent population. Under a neutral context, a vibration is detected (dashed gray line) when the population activity exceeds a criterion level (solid red line). Physical cooling of the skin (20°C) reduces afferent excitability which results in a rightward shift of the afferent response function (blue line). Separate from the physical cooling effects, the subjective experience of cold elevates the activity criterion (red dashed line) thereby raising detection thresholds in the veridical cold (dashed dark blue line) and illusory cold (dashed cyan line) conditions. (B) Distribution of simulated Pacinian mechanoreceptors activated by the threshold-level vibrations used in the psychophysical experiments. Raster plots indicate spiking patterns evoked in the simulated Pacinian afferent population for threshold stimuli in the thermoneutral (black), illusory cold (light blue), and veridical cold (dark blue) conditions. (C) Solid traces indicate frequency-response profiles for example simulated Pacinian unit when response criterion is defined as 1 spike/sec (left) or 1 spike/cycle (right). Dashed traces indicate empirical frequency-response profiles reported in Bolanowski & Verrillo (1982). For simulated and empirical data, frequency characteristics vary systematically when testing at 21°C (blue) and 29°C (red). Frequency characteristics for other Pacinian units are depicted in Figure S6. (D) Simulated detection thresholds in thermoneutral (black), illusory cold (cyan), and veridical cold (dark blue) conditions over different stimulus durations. Lines indicate average thresholds over 25 simulated populations. Shaded regions indicate s.e.m.

Prior to model fitting, we verified through simulations that, as expected, the vibration cues we tested selectively drove Pacinian afferents without recruiting slowly-adapting and rapidly-adapting afferents. Simulations further indicated that the low-amplitude vibrations tested in the staircases almost exclusively activated Pacinian receptors located on the distal finger pad (Figure 3B), in line with experimental observations that high-frequency skin vibrations decay quickly along the finger^44,79^. These results confirmed that most responsive afferents would be spatially confined within the skin region that was physically cooled in the veridical condition (Figure S5), and that vibrations did not drive receptor activity on neighboring fingers in the illusory condition.

The original TouchSim model does not simulate the effect of temperature on afferent responses. Previous empirical work has shown that cooling the skin raises response thresholds in Pacinian afferents in a frequency-dependent manner, with responses to higher frequencies more affected than those to lower frequencies^9^. To reproduce this effect in TouchSim, we varied a single parameter in the original TouchSim model (parameter 1; Methods), which controls a low-pass filter applied to the tactile input. We found that changing this parameter reproduced a number of Pacinian threshold curve features associated with temperature changes (Figure 3C; Figure S6), including the increase in absolute thresholds, the decrease in best frequencies (i.e., the frequency corresponding to maximum sensitivity), and a sharp increase in thresholds at higher frequencies with a general retention of thresholds at lower frequencies. As the original research was performed on surgically-excised Pacinian corpuscles, the findings neglect the effect of the skin tissue between the cooling device and the corpuscle, which acts as both a mechanical and temperature barrier. Indeed, the thresholds reported by Bolanowski and Verrillo^9^ were roughly an order of magnitude lower than those obtained in TouchSim or reported in the literature^40^. Nevertheless, assuming that the presence of skin tissue impacts the strength of the cooling effect but not its basic nature^9^, we treated the identified model parameter as a free parameter in our simulation that could be fitted using the psychophysically measured thresholds.

To model psychophysical detection thresholds using the simulated Pacinian activity, we implemented a simple decoder that assumes an observer detects a vibration if the total spike count in the Pacinian afferent population exceeds a specific activity level (criterion parameter). While the spiking levels in the simulated Pacinian afferents could be modulated by cold, the decoder is assumed to be naive to thermal state of the skin. To account for the subjective effect of cold perception on tactile sensitivity, we assumed that the criterion shifted (subjective parameter) when participants perceived the tested finger as feeling cold. We fitted the criterion, subjective, and TouchSim parameters on participants’ behavioral responses over all thermal and duration conditions (Methods).

Figure 3D depicts thresholds computed from model detection behavior under each condition. The model recapitulated the full pattern of threshold variations: The shorter cues and cold conditions were associated with elevated thresholds relative to thermoneutral thresholds, with the highest thresholds seen in the veridical cold condition. Overall, thresholds were elevated by 4.1 dB in the veridical and 3.3 dB in the illusory conditions compared to the thermoneutral condition, aligning with the effects observed in the psychophysical experiments.

Finally, the computational model allows us to fully dissociate physical from subjective effects by calculating expected thresholds in a hypothetical scenario where the biophysical effect of cold is present, but the subjective parameter remains unchanged from the neutral condition. This ‘afferent only’ model suggested thresholds would be elevated by a mere 0.8 dB, substantially less than the 3.3 dB elevation associated with the illusory condition, which modified the subjective parameter without the physical effect. The model therefore implies that subjective factors have a greater impact on perceptual thresholds than the purely physical effects and the contribution of physical cooling effects on thermotactile interactions is smaller than previously assumed.

## Discussion

Cold desensitization of touch is a well-established phenomenon that has been assumed to reflect biophysical impacts of cooling on signal transduction mechanisms in the skin. By leveraging thermal referral to induce an illusory cold percept on a skin region that remained physically in a thermoneutral state, our study reveals that the subjective experience of coldness alone is sufficient to desensitize touch. Moreover, behavioral and modeling results reveal that the effect of perceived cold is substantially larger than direct physical cooling effects, challenging the conventional view that thermotactile interactions originate in the somatosensory periphery.

Prior to characterizing thermotactile interactions, we first established that veridical and illusory cold percepts were similarly intense and durable over a 10-min period. Early in the adaptation period when participants experienced peak cold intensities, thermal ratings under the illusory and veridical cold conditions were statistically indistinguishable. Although participants experienced both cold conditions as less intense by the end of the adaptation period, the veridical and illusory conditions continued to be rated as colder than the thermoneutral condition. These results demonstrate the general perceptual equivalence of veridical and illusory cold.

By tracking thermal intensity over time, we quantified cold adaptation time courses and found that veridical and illusory cold intensity changed at a rate of ∼0.3°C/min, matching previously reported veridical cold adaptation rates on the forearm^11^. Adaptation of cold processing circuits was further revealed by the perceptual aftereffects observed when participants rated the intensity of the 30°C stimulus after adapting to the cold conditions: Participants rated the thermoneutral stimulus as feeling warm following cold adaptation. If thermal aftereffects resulted from adaptation of central circuits that encode thermal percepts, we would have predicted similar aftereffects following veridical and illusory cold adaptation. Because significant aftereffects were found only with veridical cold adaptation, our results are consistent with thermal adaptation being mostly confined to neural circuits encoding physical temperature rather than subjective thermal experiences^7^. The subtle aftereffects observed following 10-min exposure to the illusory cold condition potentially reflect the induction of a weak thermal referral illusion on the middle finger by warm aftereffects produced following cold adaptation of the index and ring fingers.

We found that veridical cold elevated vibration detection thresholds, consistent with earlier reports^9,21,23,25,30^. These thermal effects were independent of stimulus duration effects^28,45,46^. Extending on prior findings, we discovered that the illusory experience of cold – induced by thermal referral^4–7,30^ – similarly reduces tactile sensitivity. A role for subjective cold experience was further supported by establishing a relationship between changes in tactile thresholds and cold intensity ratings, which declined gradually with adaptation despite the physical temperature of the cooled fingers remaining a constant 20°C. Because the middle finger was maintained at 30°C to induce thermal referral, we considered whether cold stimulation of the flanking fingers or the perception of illusory cold might have triggered vasoconstriction on the middle finger to impact tactile signaling. Such a physiological response, which occurs when the body and extremities are cooled^31^, could reflect an adaptive thermoregulatory process that minimizes heat loss and preserves core temperature^33^, analogous to reflexive pupil constriction elicited by both real and illusory brightness^47^. However, thermal referral was not associated with changes in the temperature of the middle finger, ruling out indirect vasoconstriction. These findings reveal that thermotactile interactions result from both physical and subjective factors, with the latter likely involving central mechanisms.

How does skin cooling modulate tactile sensitivity? Using a neurophysiologically grounded model of somatosensory afferents^8^, we found that our vibration stimuli selectively drove activity in Pacinian afferents. To account for the physical effects of skin cooling, we allowed the spiking properties of these afferents to vary with skin temperature, controlled by a single model parameter (Methods). By fitting this parameter to participants’ detection thresholds, together with parameters defining the activity levels required for detection under thermoneutral and cold states, the model reproduced the full range of observed behavioral patterns. Extending beyond the tested conditions, we simulated afferent responses over an expanded range of vibration frequencies and replicated cooling effects on Pacinian response profiles^9^, including reductions in absolute sensitivity and tuning curve shifts. Although model thresholds differed in magnitude from empirical thresholds, these differences could be attributed to the fact that the recordings were obtained from isolated Pacinian corpuscles rather than mechanoreceptors embedded in a temperature- and vibration-filtering skin matrix. These results are nevertheless consistent with the prevailing view that cold attenuation of vibration sensing can be attributed, in part, to temperature-mediated changes to Pacinian afferent sensitivity.

Our novel finding – the elevation of detection thresholds by illusory cold without concomitant changes in skin temperature – reveals the modulatory influence of the subjective experience of cold on tactile sensitivity. To model detection behavior, we defined a threshold level of activity in the Pacinian afferent population above which a stimulus was considered detected. This threshold is conceptually related to the criterion parameter in signal detection theory^48^ or to a decision boundary in bounded evidence accumulation models^49^, both of which may be linked to activity in frontal and parietal brain regions^49–51^. According to our model, the experience of cold elevates this threshold, thereby requiring stronger vibrations to evoke the spiking levels needed for detection. An alternative, but functionally equivalent, model is that cold-related activity reduces the effective gain of vibrations responses via divisive normalization^52^. In this framework, cold-related activity increases the normalization pool activity which proportionally reduces the gain of vibration-signaling neurons downstream of the Pacinian afferents. Divisive normalization has been proposed as a canonical computation underlying contextual modulation of touch^53^ and has been linked to activity in somatosensory cortex^54^. Consistent with this view, recent neurophysiological studies demonstrate the convergence of thermal and tactile signals in somatosensory cortex^55,56^.

However, gain modulation of tactile responses could occur at multiple levels along the somatosensory neuroaxis spanning spinal cord^57^, dorsal column nuclei^58^, and the ventrobasal thalamic complex^59^. Because tactile threshold changes related to subjective cold intensity rather than the physical temperature of the skin, the critical neural signals modulating touch must originate in a substrate that encodes subjective cold experiences. Given the established relationship between posterior insula cortex and cold sensing^60–65^, this association area is a strong candidate for providing the modulatory thermal signals. Additional neurophysiological studies are needed to determine whether the perceptual impact of veridical or illusory cold on touch results from criterion-shifting, gain control, or some other computation, in addition to establishing their neural basis.

Our findings demonstrate that the mere experience of cold is sufficient to attenuate tactile sensitivity, even in the absence of actual skin cooling. This result reveals that thermotactile interactions are not solely determined by peripheral biophysical mechanisms but can arise from central processes that reflect sensory expectations and prior associations. Conceivably, the illusory cold percept produced by thermal referral reactivates neural circuits that represent the association between cold sensations and reduced tactile processing, learned through repeated co-occurrence of physical cooling and diminished afferent drive. Such circuits may serve an adaptive function in down-weighting touch when sensory evidence is degraded and recruiting alternative behavioral strategies^66^, thereby minimizing inference errors and conserving metabolic resources. Indeed, because thermal modulation of skin properties and somatosensory signaling occurs faster than one can experience the thermal cues^67^, a rational observer may assume that touch is altered any time a thermal stimulus is experienced. Finally, given that thermal perception is closely tied to body ownership^68^, tactile sensitivity changes may relate to rapid alterations in body schema such as those seen with the Rubber Hand Illusion^69–71^. Regardless of the exact mechanism, the subjective experience of cold appears to serve as a predictive signal that recruits modulatory pathways capable of scaling tactile gain, analogous to how expectation-based mechanisms elicit genuine physiological responses related to pain^72,73^, pleasure^74^, and even immune system function^75^. Thermotactile interactions thus highlight how multimodal experiences and expectations shape the neural computations underlying somatosensation. If such expectations can be recalibrated through novel multimodal experiences, they may offer a new avenue for interventions aimed at restoring impaired tactile sensitivity.

## Acknowledgements

This work was supported by grants from the National Institutes of Health (R01NS127777, F31NS135914) and BCM Seed Funds. JV was supported by the Secretaría de Educación, Ciencia, Tecnología e Innovación de la Ciudad de México (Grant Number: SECTEI/103/2022). We thank Jian Chen and Jing Lin for their contributions in programming and device engineering, Katie Steck for helping with the artwork in Figure 1, David Lipshutz and Javier Medina for helpful discussions, and Joseph Banerjee for helping set up the simulation environment.

## Author contributions

JR, JV, and JMY conceived experiments. JR performed the experiments. HPS implemented the model. JR, JV, BK, and HPS performed the data analysis. JR and JMY wrote the manuscript with input from all authors.

## STAR Methods

### PARTICIPANTS

Thirteen subjects (7 females; ages 24–38, mean = 29.44, SD = 4.16 years) participated in both the adaptation and threshold experiments. Participants completed the 2 experiments on separate days (inter-session-interval = 72.5 days). Eight out of 13 participants were right hand dominant (Edinburgh Handedness score^76^: 60.55±25.68). No participants reported a history of neurological disorders, psychiatric conditions, sensory impairments, or intolerance to thermal stimuli. Participants provided written informed consent and received monetary compensation. Experimental procedures were approved by the Institutional Review Board at Baylor College of Medicine, and conducted in accordance with the Declaration of Helsinki and the Belmont Report.

## METHODS DETAILS

### Thermotactile stimulation device

Participants experienced thermal and mechanical stimulation through a custom thermotactile stimulation system^12^. Mechanical (vibrotactile) cues were delivered only to the middle finger while thermal stimulation was delivered independently to the index, middle, and ring fingers. The distal finger pad of each finger contacted a separate rectangular thermo-electric Peltier element (24.6mm × 12.3mm; 9W, Laird Thermal Systems). The Peltier contacting the middle finger was mounted on a shaker motor (LTV Ling Altec) that was controlled via custom software on a PC (Intel) and external amplifier (Siglent SPA1010). Shaker motor performance was recorded using an accelerometer (Kistler 8786A5). All stimulator components were mounted to an aluminum frame (80/20, McMaster-Carr). Participants sat comfortably with their left arm supported in a supinated position and their left hand stably positioned beneath the thermo-electric devices. Participants wore noise-attenuating earmuffs (3M Peltor) to mask sounds produced by the shaker motor. The Peltier contacting the middle finger was lowered into the distal pad 0.5mm beyond initial contact. System performance for delivering thermal stimulation (Figure S1) and mechanical stimulation at 250Hz has been described previously^12^.

### Temperature recording hardware

The temperature of the Peltiers and skin were recorded using thermistors (Ultra Thin 10k Ohm Thermistor B395, Adafruit Industries). Contact-less skin temperature measurements were also recorded using an infrared thermal camera (MLX90640, 110° POV, Adafruit Industries, Thermal Camera Breakout Circuit) in conjunction with a microcontroller (Teensy 4.0).

### Electrocardiogram (ECG) recording hardware

To record real-time cardiac data during the detection threshold experiment, we used an Single Lead Heart Rate Monitor (AD8232, Adafruit Industries) via an Arduino (UNOv3, Arduino) in conjunction with a 3-lead cable and snap-on EKG Monitoring Electrodes (MedGel) and a data acquisition system (AlphaLab SNR, AlphaOmega). Three disposable gel electrodes were placed on the chest of each participant approximately 5 inches below the clavicle, arranged according to Eintoven’s Triangle^77^.

### Thermal intensity rating task

The purpose of this experiment was to quantify the quality (cool vs warm) and intensity of participants’ thermal percepts under sustained exposure to the thermoneutral (30°C across fingers), veridical cold (20°C across fingers), and illusory cold (20°C on the index and ring fingers with 30°C on the middle finger) conditions. Participants reported the perceived thermal intensity on the middle finger using a color-coded Visual Analog Scale (VAS) that ranged continuously (1920 levels) from dark blue (representing maximum innocuous cold) to dark red (representing maximum innocuous hot). Participants indicated the color along the bar corresponding to their subjective thermal rating by mouse click. Prior to testing, participants were calibrated to the VAS by delivering 20°C, 30°C, and 40°C (uniform over the fingers) and explicitly associating these levels to dark blue, white, and dark red, respectively. In separate 10-min adaptation periods, participants experienced the neutral and 2 cold conditions and responded every 20 seconds with visual prompts. After each adaptation period, participants underwent a 10-min readaptation period comprising uniform 30°C stimulation over the 3 fingers. Participants were also prompted to respond using the VAS at 20-s intervals during the readaptation periods.

### Vibration detection task

The purpose of this experiment was to quantify each participant’s detection thresholds under 3 different thermal conditions (neutral, veridical cold, illusory cold) and 2 vibration durations (100ms, 500ms). Vibration detection thresholds were determined using a 2-interval, 2-alternative forced-choice paradigm. On each trial (inter-trial interval = 500-1500ms), a 250-Hz vibration was delivered to the middle finger in 1 of 2 stimulus intervals and participants reported by button press whether they experienced the test stimulus in the first or second interval. The stimulus-containing interval was randomized across trials and the time between stimulus intervals varied randomly from 500-1500ms to prevent participants from anticipating stimulus delivery. A 1-Up/2-Down adaptive staircase determined vibration amplitudes: If participants reported detecting the stimulus in the incorrect interval, stimulus amplitude on the subsequent trial was increased. Conversely, if participants responded correctly on 2 trials, stimulus amplitude on the subsequent trial was decreased. Amplitudes changed by a factor of 2 prior to the 5^th^ reversal and by a factor of 4 subsequently with the staircase terminating after 10 total reversals. Three staircases (2 descending, 1 ascending) were performed serially for each condition within a 10-min period. The descending and ascending staircases started at 3mm and 0.01mm, respectively. Conditions were tested in a counterbalanced, randomized order across participants. Each testing period was preceded by a 10-min adaptation period to 30°C uniform stimulation over fingers.

## QUANTIFICATION AND STATISTICAL ANALYSIS

Data were assessed for normality using the Shapiro-Wilk test. Unless otherwise noted, group-level parametric tests comprised repeated-measures ANOVA and post-hoc comparisons while nonparametric tests comprised Friedman’s test and post-hoc Wilcoxon signed-rank comparisons. All post-hoc tests were corrected for multiple comparisons. Analyses were conducted in MATLAB R2023a and RStudio.

### Thermal intensity ratings

Thermal intensity ratings recorded during the adaptation and readaptation periods were normalized by each participant’s maximum unsigned intensity rating. Accordingly, a value of −1 corresponded to the participant’s maximally cold rating while a value of 1 corresponded to the participant’s maximally warm rating. Normalized intensity ratings were averaged over the first and last minute of each adaptation and readaptation period. Adaptation rates (change in intensity as a function of time) were determined by linear regression. In group-level tests, we compared intensity ratings and adaptation rates (slope parameters) across thermal conditions.

### Vibration detection thresholds

A threshold was computed for each staircase according to the mean amplitude over the final 5 staircase trials. For each participant, thresholds across conditions (thermal and duration) were normalized by converting threshold values to z-scores. In group-level tests, we evaluated whether thresholds modulated according to thermal condition (3 levels), duration (2 levels), and staircase (3 levels) using aligned rank transform (ART) ANOVA. Post-hoc comparisons used estimated marginal means and permutation tests.

### Relating detection thresholds to thermal intensity ratings

We first computed each participant’s average intensity ratings during the adaptation periods corresponding to the times during which participants tended to perform the final 5 staircase trials, namely from (min:sec) 2:40-3:20, 5:40-6:20, and 9:00-10:00. This yielded a “staircase-tied” thermal intensity rating for each staircase under each thermal condition. To evaluate covariation between detection thresholds and thermal intensity ratings for each participant, we computed the Spearman’s correlation between the staircase-tied thermal intensity ratings and normalized thresholds for each staircase (z-scored within durations). In a group-level test, we determined whether the group-averaged correlation differed significantly from 0 using a one-sample t-test.

### Cardiac data analysis

EKG data were sampled at 44kHz and preprocessed using a zero-phase Butterworth bandpass filter (0.5–45 Hz, 4th order) implemented in MATLAB (butter, filtfilt). The lower cutoff (0.5 Hz) removed baseline wander and slow drift, while the upper cutoff (45 Hz) attenuated muscle and high-frequency electrical noise. Forward–backward filtering preserved phase information critical for R-peak alignment. First, R-peaks were detected from the preprocessed EKG, and successive R–R intervals were computed for each of the ∼3min recordings per subject (18 recordings per subject). Each cardiac cycle was then divided into two phases based on the duration of the R–R interval. Systole was defined as the first one-third of the interval following each R-peak to approximately coincide with the end of the T wave, and diastole was defined as the remaining two-thirds.

For each participant, we took the amplitudes of the last 10 trials for each adaptive staircase, which yielded 30 trials per temperature condition. Each trial was assigned a cardiac phase by comparing the vibration onset to the time window that corresponded to systole or diastole. If the vibration began within the first 1/3 of the R–R interval, the trial was labeled as systole; otherwise, it was labeled as diastole. Because sample sizes were unequal across conditions, we modeled trial-level vibration amplitude with a linear mixed-effects model (Gaussian identity link) including cardiac phase (diastole, systole), duration (100, 500ms), and temperature (neutral, illusory cold, veridical cold) as fixed effects, and a random intercept for subject to account for repeated measures. We fitted the model using restricted maximum likelihood (REML) in MATLAB (fitlme). Significance of fixed effects was assessed using Satterthwaite’s approximation to degrees of freedom. Statistical significance was set at α = 0.05.

### Skin temperature analysis

To determine whether manipulating the temperature of the index and ring fingers (D2 and D4) modulated the temperature of the unloaded middle finger, we implemented a protocol comprising three contiguous 10-min epochs: (i) 30°C application to D2 and D4 (pre-thermal referral), (ii) 20°C application to D2 and D4 (thermal referral induction), and (iii) 30°C application to D2 and D4 (post-thermal referral). Independent temperature measurements were recorded using a thermistor and infrared camera. We acquired infrared thermal images of the skin at 2Hz. The raw 32×24 pixels were interpolated to generate a 1150×1550 grid. Within this grid, we defined a 400×400 mm pixel region of interest (ROI) (∼1.25 cm^2^) centered over the distal pad of the middle finger. For each participant, we extracted the mean temperature (°C) within the ROI for each frame. We conducted a repeated-measures ANOVA to determine if the average middle finger temperature varied significantly according to the 3 epochs. For all tests, statistical significance was set at α = 0.05.

The infrared camera was used similarly to quantify the temperature of the middle finger pad on the middle finger when the distal pad contacted the Peltier set to 20°C. To determine the temperature of skin contacting the Peltier, when the view of the infrared camera was occluded, a thermistor was attached directly to the skin.

## AFFERENT AND BEHAVIOR MODELING

To model physical and subjective factors associated with cold modulation of vibration detection thresholds, we implemented a mechanistic model to simulate perceptual stimulus detection.

To account for physical effects, we simulated peripheral afferent activity using TouchSim^8^, a computational model that recreates neural spiking responses to tactile stimuli. In initial simulations, we confirmed that our high-frequency, low-amplitude vibration stimuli did not excite slow-adapting type 1 or rapidly-adapting fibers, in line with their widely reported neurophysiological response properties^78^. As expected, Pacinian fibers were activated by these stimuli and our model therefore focused on this afferent class only. Vibrotactile stimuli elicit skin oscillations that propagate through the skin and can therefore elicit responses in receptors located some distance away from the contact point^44,79^. In our simulations, we found activity along the length of the stimulated (middle) finger in some conditions, but this never reached the palm or other areas of the hand. We therefore focused our simulation on the middle finger only, excluding the rest of the hand. While PC response properties are very stereotyped, there is some individual variability in overall sensitivity and threshold curves^80^; moreover, their precise number and location within the skin will vary somewhat across participants. To account for such variability, we resampled a new PC population 25 times, with afferents terminating at random locations on the finger according to their estimated innervation densities^81^ and randomly assigning 1 of 4 PC models available in TouchSim.

Previous empirical work has shown that cooling the skin raises response thresholds in PC afferents in a frequency-dependent manner, with higher frequencies more affected than lower frequencies^9^. To reproduce this effect in TouchSim, we varied a single parameter in the original TouchSim model (parameter 1), which controls a low-pass filter applied to the tactile input. As the original research was performed on surgically-excised Pacinian corpuscles, the findings neglect the effect of the skin tissue between the cooling device and the corpuscle, which will act as both a mechanical and temperature barrier. Indeed, the thresholds reported by Bolanowski and Verrillo (1982) were roughly an order of magnitude lower than those obtained in TouchSim or reported in the literature. Accordingly, fitting of the TouchSim parameter to account for cooling was performed only after the empirical (29°C) curves^9^ were rescaled to match the threshold curves produced by the original TouchSim. Beyond impacting receptor sensitivity, neglecting the presence of skin tissue likely overestimates the effect of temperature by removing the tissue’s insulating effect.

To simulate detection of the vibrotactile stimulus, we assumed a central criterion on spiking activity, with successful detection once this criterion was exceeded. Prior research has shown that individual action potentials in PC fibers are not perceptually detectable, while higher stimulation frequencies are^82^, supporting the idea of a central threshold. In line with these findings, we set the total population spiking level across all simulated PC afferents as the criterion, which was then fit to the empirical behavioral data. While more complex criteria are possible, the simple population spike count employed here reproduced the observed effect that stimuli with longer durations require lower amplitudes to reach threshold and therefore was sufficient to capture this effect.

To match our model against the psychophysical threshold data and test whether physical or subjective factors could best account for the observed response patterns, we fitted a 3-parameter model to the psychophysical data. The free parameters were the criterion parameter (reflecting the activity level required for detection in the neutral condition), a subjective parameter (capturing a shift of the criterion in the illusory and veridical conditions), and finally a modified TouchSim parameter (capturing thermal effects on PC spiking), assumed to be active in the veridical cold condition. The cost function was the sum of the squared errors in all six experiments (3 conditions x 2 durations), expressed as the logarithm of the ratio between simulated and average measured perceptual thresholds. The logarithm was chosen to account for the fact that thresholds can vary over orders of magnitude. Optimization was performed using the *pybads* package^83^, a Python implementation of Bayesian Adaptive Direct Search^84^ that is suitable for fitting multiple parameters simultaneously in non-convex derivative-free optimization problems. Optimization runs converged for all 25 sampled population models.

## Data and code availability

All relevant datasets and analysis scripts generated during this study are available upon request from the corresponding author.

## Ethics statement

This research was performed with approval from Baylor College of Medicine’s Institutional Review Board (IRB; H-54490), consistent with ethical guidelines outlined in the Declaration of Helsinki and Belmont Report.

---

**Figure S1.**
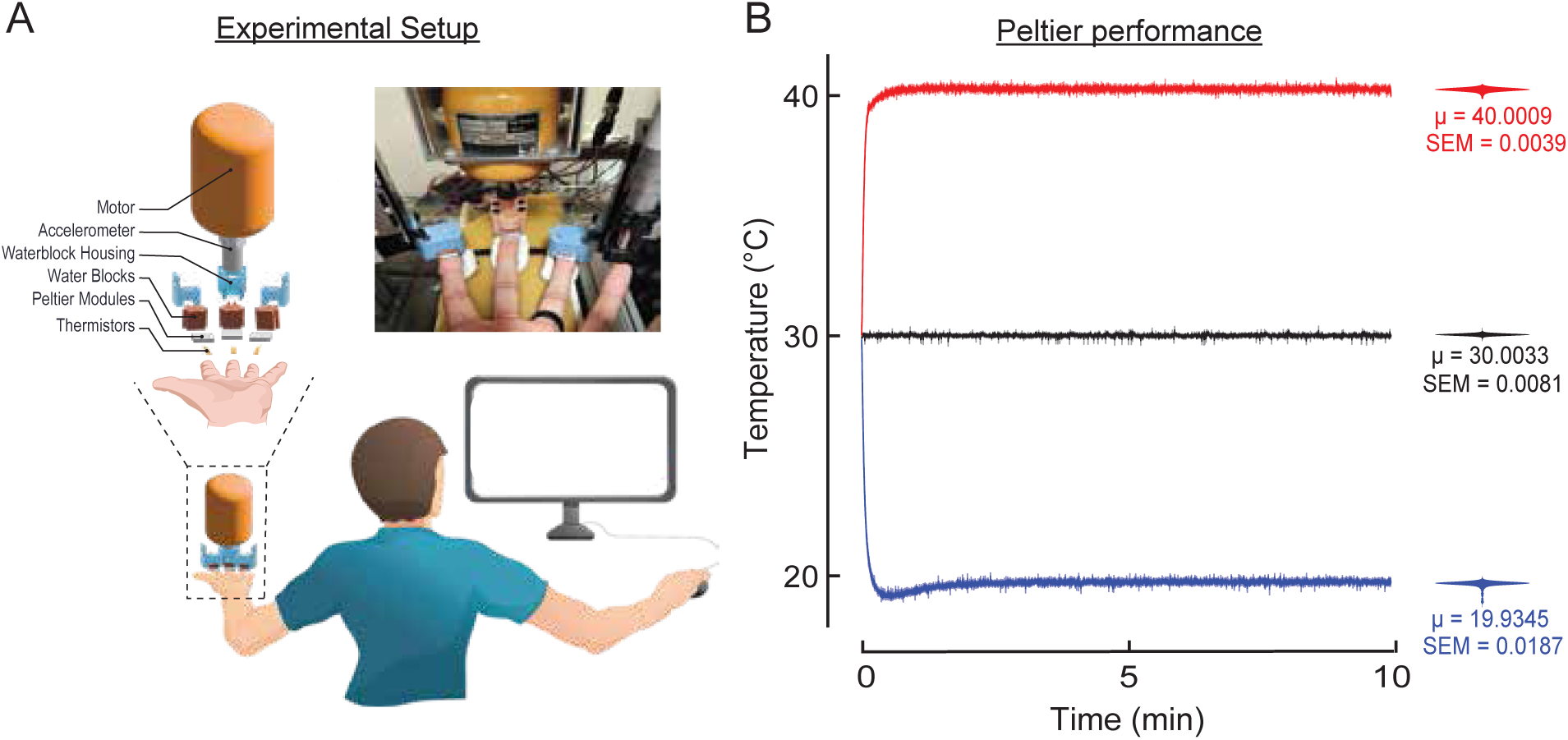
Experimental setup. A) Schematic of the thermotactile stimulation device. Participants placed the index, middle, and ring fingers of their left hand on 3 independently controlled Peltier elements. The middle finger Peltier was mounted to a shaker motor to deliver controlled vibrotactile stimuli. The inset shows individual device components. Participants viewed instructions, timing cues, and response prompts on a computer monitor. Participants responded using mouse clicks. (B) Peltier performance for delivering cold, neutral, and hot temperatures. Traces indicate mean time course (5 repetitions) for 40°C (red), 30°C (black), and 20°C (blue) over 10 min under thermal load (fingertip). Violin plots indicate the mean (μ) and SEM of temperatures over the steady state (the point at which the desired temperature is first reached until the end of 10 min). See Ramirez et al., 2025 for full description.

**Figure S2.**
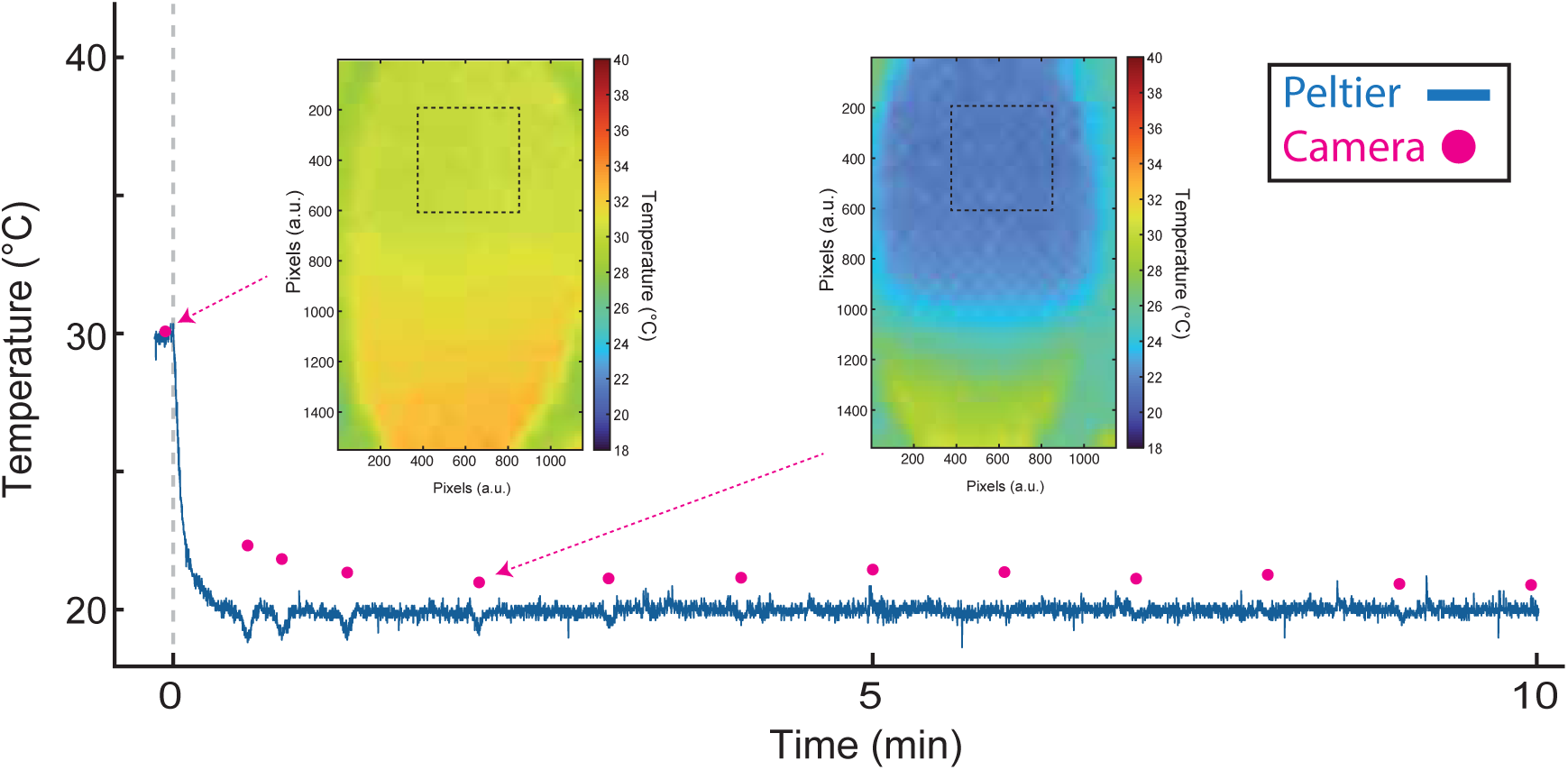
Skin temperature profile when cooled by Peltier device set to 20°C. Red dots indicate surface temperature of the middle fingertip measured using contactless thermal imaging while the Peltier device was initially held at 30°C and then reduced to 20°C for 10 minutes beginning at time = 0. Because the Peltier device occluded the camera’s view of the contacted distal finger pad, contactless measurements were recorded by retracting the middle finger from beneath the Peltier element briefly after 15s, after 30s, and at 1-min intervals thereafter over the 10 minute period. Inset plots depict thermal images of the distal finger pad acquired prior and during active cooling. The dashed squares indicate the region-of-interest over which temperature was computed during each sample. The blue trace indicates temperatures recorded continuously from a thermistor attached to the distal finger pad. Note that the thermistor was in physical contact with the Peltier when the distal finger pad was positioned beneath directly beneath the cooling element. Both contactless and physical thermal measurements indicate the distal finger pad rapidly cooled to ∼20°C and remained cool for the duration of recording.

**Figure S3.**
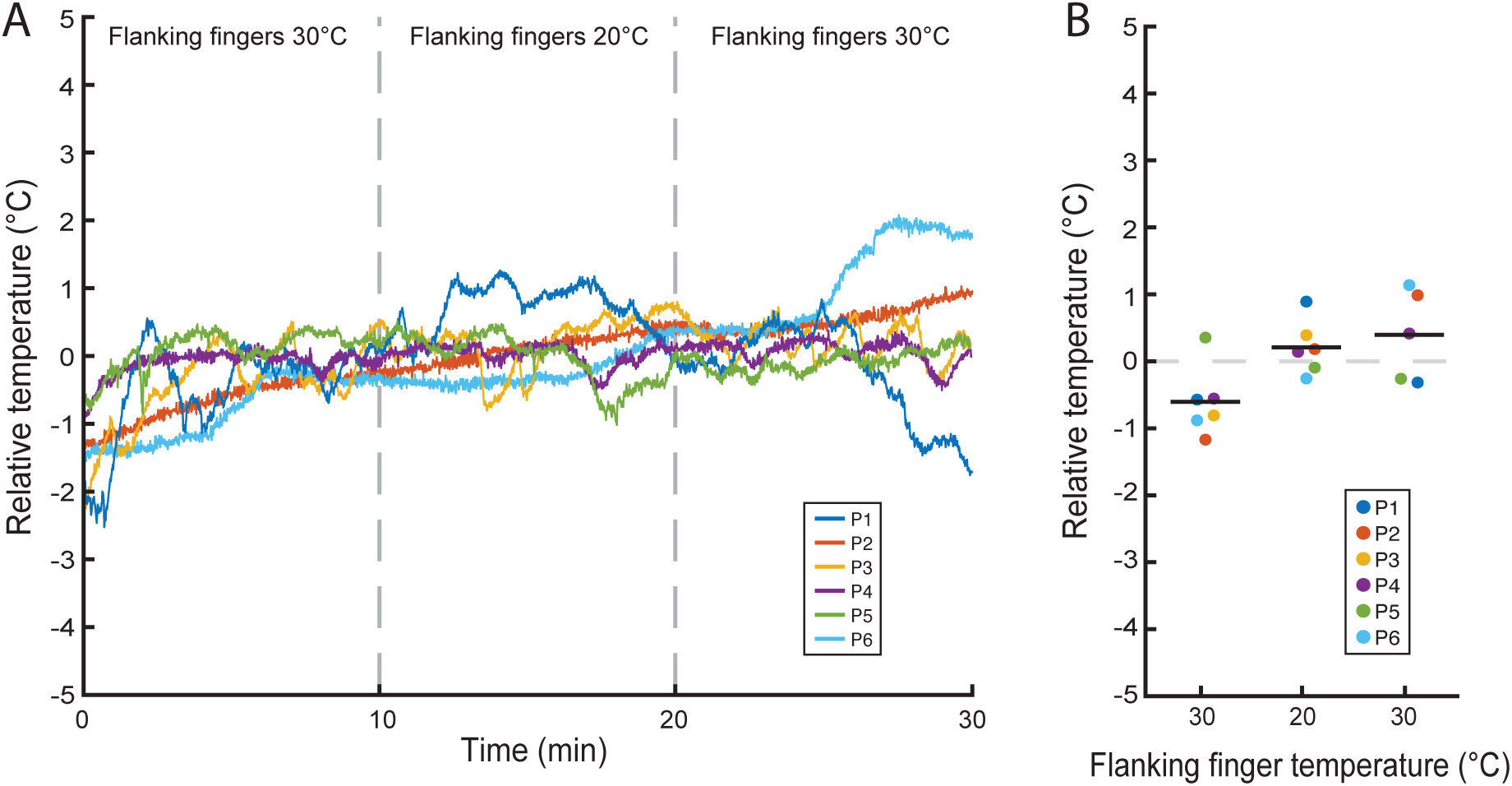
Distal pad of middle finger does not cool when flanking fingers experience cold stimulation. (N = 6). The temperature of the <u>unloaded</u> distal finger pad on the middle finger was measured using a thermistor when flanking index and ring fingers were maintained at 30°C (pre-thermal referral), cooled to 20°C (thermal referral induction), and returned to 30°C (post-thermal referral). A) Traces indicate temperature changes relative to the mean temperature computed over the full 30-min interval within individual participants. B) Markers indicate each participant’s average temperature changes (relative to mean temperatures) computed for each 10-min epoch. Horizontal lines indicate group average. Despite a trend for finger warming over the 3 epochs, finger temperature did not vary significantly according to epoch (F(2,10) = 6.7×10^-3^, p = 0.94). Cooling of the middle finger would be expected if cold-stimulation of the flanking fingers induced indirect vasoconstriction.

**Figure S4.**
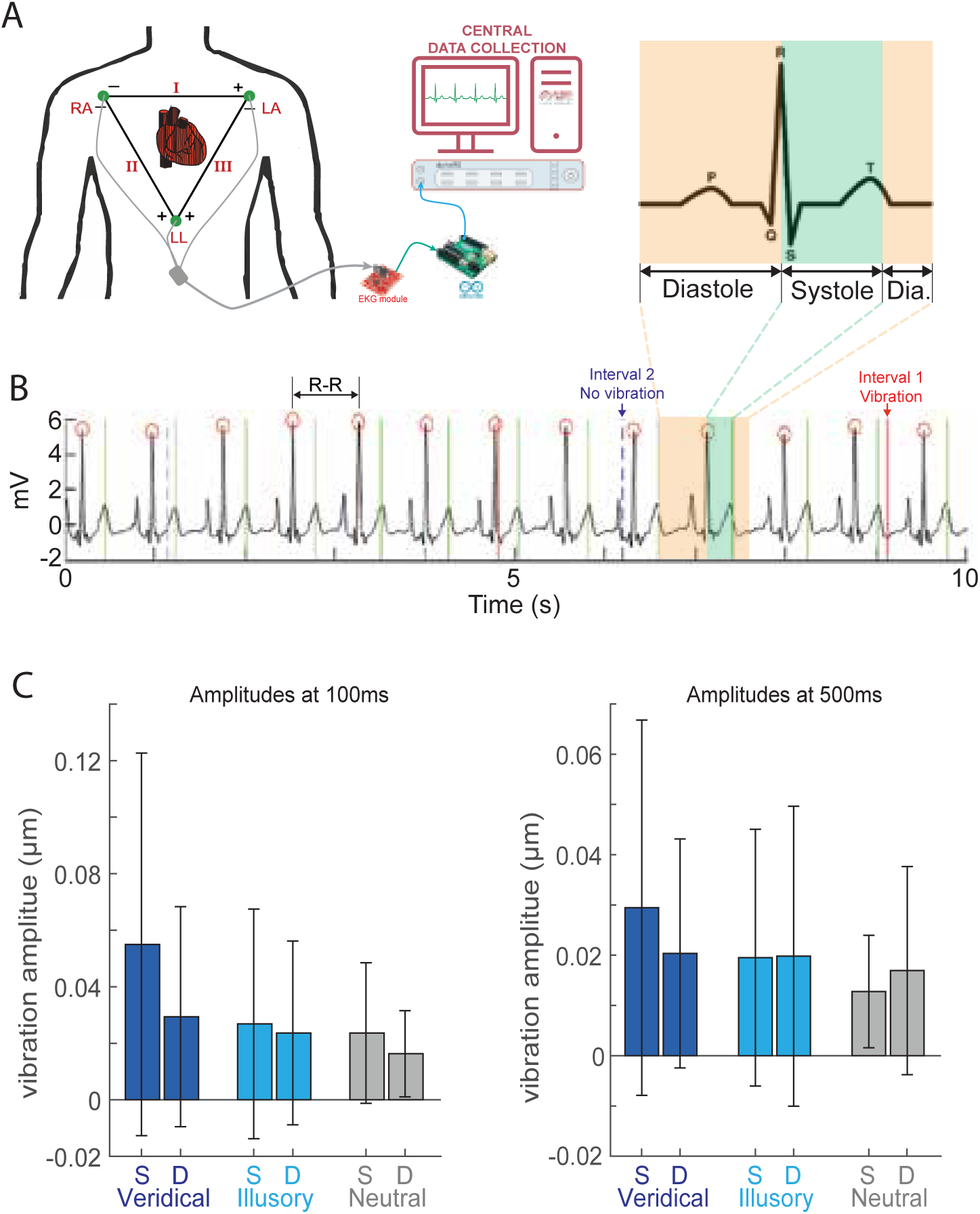
Relationship between stimulus amplitude and cardiac phase. A) Diagram (*left*) indicates EKG electrode placement. Standard PQRST electrocardiogram of one heartbeat cycle (*right*) is color-coded to indicate systolic (green) and diastolic (orange) windows. B) 10-sec EKG recording sample from example participant. R-peaks are outlined with red circles. The systolic phase is defined as 1/3 the R-R interval. Solid red lines indicate stimulus intervals containing a vibration while dashed blue lines indicate stimulus intervals without vibrations. C) Tested vibration amplitudes (over last 10 trials of each staircase) sorted by temperature condition and cardiac phase (‘S’ = Systolic, ‘D’ = Diastolic) for 100-ms (left) and 500-ms (right) vibrations (n=13). Bar plots indicate average stimulus amplitudes in each condition. Errorbars indicate s.e.m. Vibration amplitudes differed significantly according to thermal condition (main effect; F = 7.71, p = 4.83×10^-4^) and duration (main effect; F = 9.19, p = 2.52×10^-3^). No main and interaction effects involving cardiac phase achieved statistical significance (all ps > 0.05).

**Figure S5.**
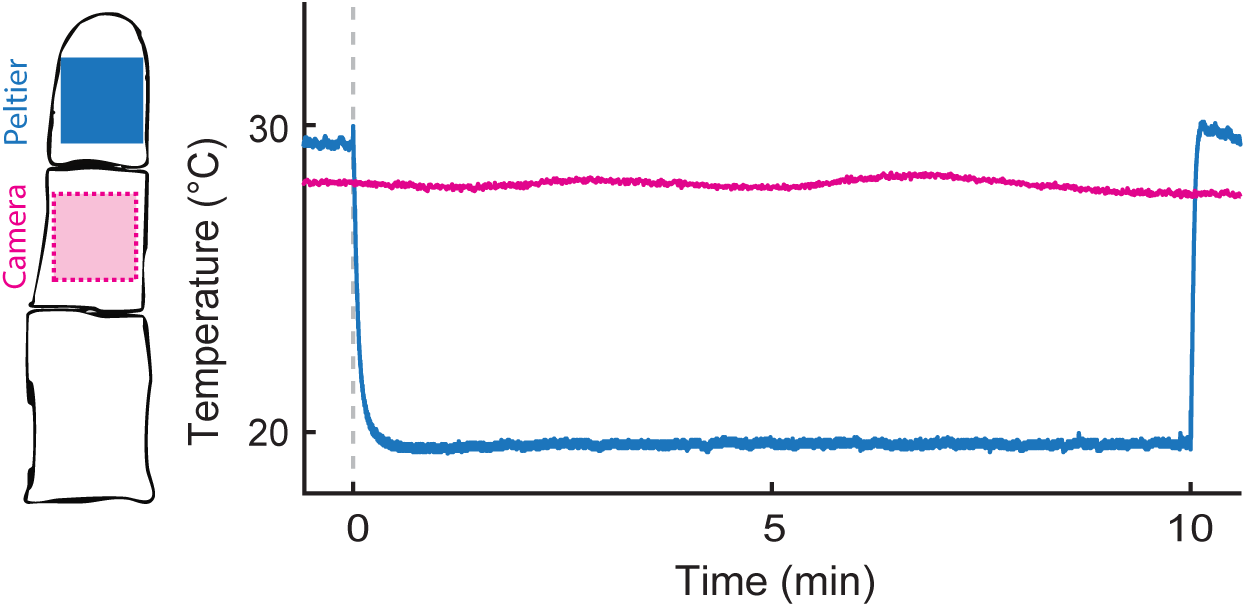
Middle finger pad temperature is unchanged when the distal finger pad experiences cold stimulation. Blue trace indicates thermal recordings from the thermistor on the Peltier element (blue square) contacting the distal finger pad. Magenta trace indicates contactless temperature measured over a region-of-interest (magenta square; ∼12.7mm^2^) centered on the middle finger pad. Continuous optical thermal recordings from the middle finger pad were sampled at 2Hz. These data reveal that physical cooling of the distal finger pad does not cause direct or indirect cooling of the middle finger pad.

**Figure S6.**
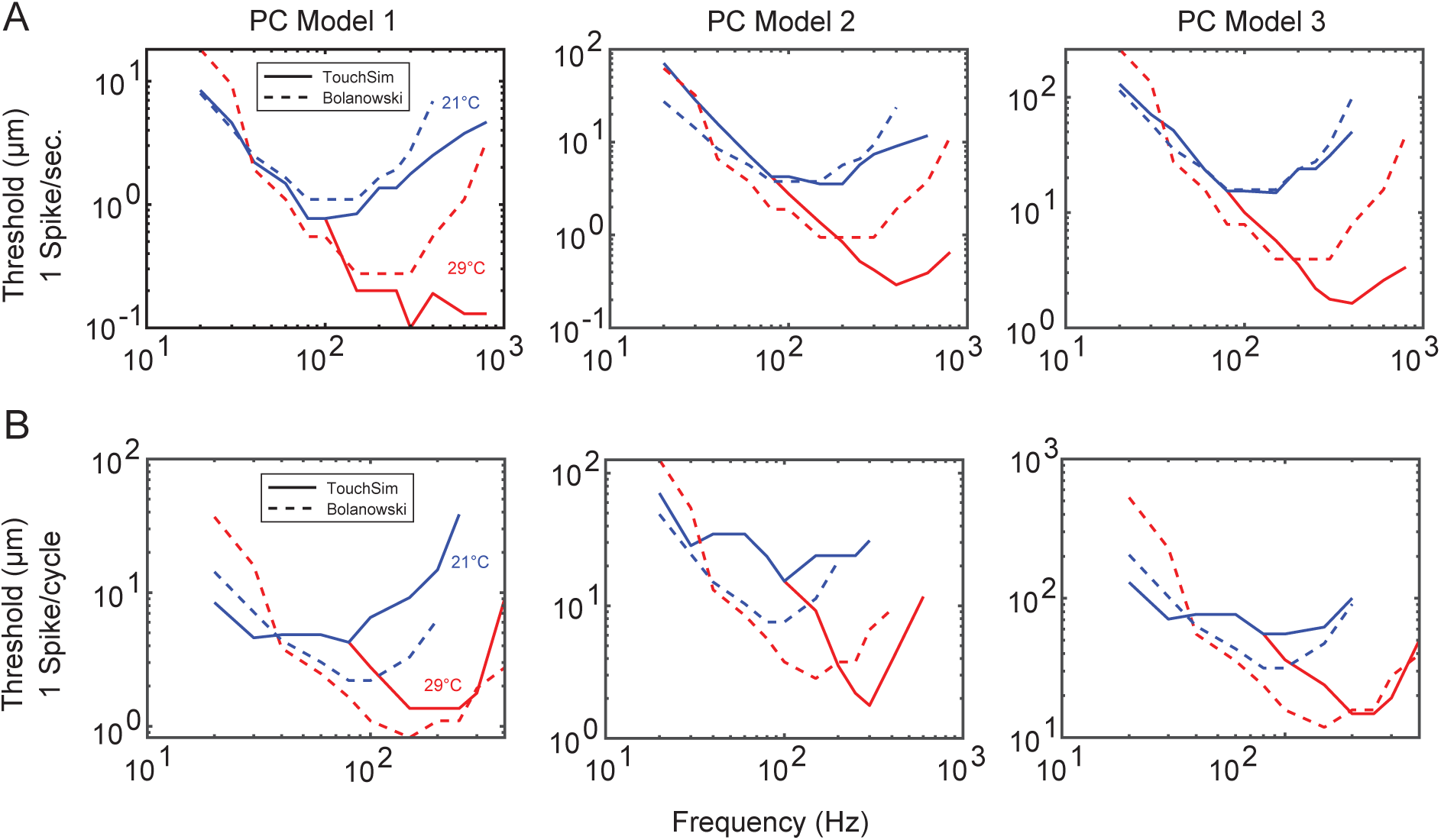
Empirical and simulated Pacinian afferent threshold functions. Traces indicate thresholds defined as the stimulus amplitude required to evoke A) 1 spike per second or B) 1 spike per stimulus cycle as a function of vibration frequency. Solid traces show response curves for 3 simulated Pacinian afferents (TouchSim, Saal et al., 2017) and dashed traces show empirical response curves from Pacinian afferent recording (Bolanowski & Verrillo, 1982). Colors indicate response curves determined under 21°C (blue) and 29°C (red). In empirical and simulated data, cooling elevates afferent thresholds over higher frequency ranges. While PC response properties are very stereotyped, there is some individual variability in overall sensitivity and threshold curves (Muniak et al., 2007); moreover, their precise number and location within the skin will vary somewhat across participants. To account for such variability, we resampled a new PC population 25 times, with afferents terminating at random locations on the finger according to their estimated innervation densities (Johansson & Vallbo, 1979) and randomly assigning 1 of 4 PC models available in TouchSim (3 depicted here and additional PC model depicted in Figure 3C).

## References

1. Ishiko, N., and Loewenstein, W.R. (1961). Effects of temperature on the generator and action potentials of a sense organ. J. Gen. Physiol. 45, 105–124. 10.1085/JGP.45.1.105.

2. de Lange, F.P., Heilbron, M., and Kok, P. (2018). How Do Expectations Shape Perception? Trends Cogn. Sci. 22, 764–779. 10.1016/J.TICS.2018.06.002.

3. Green, B.G. (1977). Localization of thermal sensation: An illusion and synthetic heat. Percept. Psychophys. 22, 331–337. 10.3758/BF03199698/METRICS.

4. Cataldo, A., Ferrè, E.R., Di Pellegrino, G., and Haggard, P. (2016). Thermal referral: evidence for a thermoceptive uniformity illusion without touch. Sci. Rep. 6. 10.1038/SREP35286.

5. Ho, H.N., Watanabe, J., Ando, H., and Kashino, M. (2011). Mechanisms underlying referral of thermal sensations to sites of tactile stimulation. J. Neurosci. 31, 208–213. 10.1523/JNEUROSCI.2640-10.2011.

6. Ho, H.N., Watanabe, J., Ando, H., and Kashino, M. (2010). Somatotopic or spatiotopic? Frame of reference for localizing thermal sensations under thermo-tactile interactions. Atten. Percept. Psychophys. 72, 1666–1675. 10.3758/APP.72.6.1666.

7. Ho, H.N., Chow, H.M., Tsunokake, S., and Roseboom, W. (2019). Thermal-Tactile Integration in Object Temperature Perception. IEEE Trans. Haptics 12, 594–603. 10.1109/TOH.2019.2894153.

8. Saal, H.P., Delhaye, B.P., Rayhaun, B.C., and Bensmaia, S.J. (2017). Simulating tactile signals from the whole hand with millisecond precision. Proc. Natl. Acad. Sci. U. S. A. 114, E5693–E5702. 10.1073/PNAS.1704856114/SUPPL_FILE/PNAS.1704856114.SM02.AVI.

9. Bolanowski, S.J., and Verrillo, R.T. (1982). Temperature and criterion effects in a somatosensory subsystem: a neurophysiological and psychophysical study. J. Neurophysiol. 48, 836–855. 10.1152/JN.1982.48.3.836.

10. Filingeri, D., Zhang, H., and Arens, E.A. (2017). Characteristics of the local cutaneous sensory thermoneutral zone. J. Neurophysiol. 117, 1797. 10.1152/JN.00845.2016.

11. Kenshalo, D.R., and Scott, H.A. (1966). Temporal course of thermal adaptation. Science 151, 1095–1096. 10.1126/SCIENCE.151.3714.1095.

12. Ramirez, J.C., Vergara, J., Lin, J., Chen, J., and Yau, J.M. (2025). A novel device for studying temperature and touch interactions. J. Neurosci. Methods 423. 10.1016/J.JNEUMETH.2025.110547.

13. Weber, A.I., Krishnamurthy, K., and Fairhall, A.L. (2019). Coding Principles in Adaptation. Annu. Rev. Vis. Sci. 5, 427–449. 10.1146/ANNUREV-VISION-091718-014818.

14. Whitmire, C.J., and Stanley, G.B. (2016). Rapid Sensory Adaptation Redux: A Circuit Perspective. Neuron 92, 298–315. 10.1016/J.NEURON.2016.09.046.

15. Fairhall, A.L., Lewen, G.D., Bialek, W., and De Ruyter van Steveninck, R.R. (2001). Efficiency and ambiguity in an adaptive neural code. Nature 412, 787–792. 10.1038/35090500.

16. Schwartz, O., Hsu, A., and Dayan, P. (2007). Space and time in visual context. Nat. Rev. Neurosci. 8, 522–535. 10.1038/NRN2155.

17. Seriès, P., Stocker, A.A., and Simoncelli, E.P. (2009). Is the homunculus “aware” of sensory adaptation? Neural Comput. 21, 3271–3304. 10.1162/NECO.2009.09-08-869.

18. Verrillo, R.T., and Bolanowski, S.J. (2003). Effects of temperature on the subjective magnitude of vibration. Somatosens. Mot. Res. 20, 133–137. 10.1080/089902203100105163.

19. Verrillo, R.T., and Bolanowski, S.J. (1986). The effects of skin temperature on the psychophysical responses to vibration on glabrous and hairy skin. J. Acoust. Soc. Am. 80, 528–532. 10.1121/1.394047.

20. Gescheider, G.A., Thorpe, J.M., Goodarz, J., and Bolanowski, S.J. (1997). The effects of skin temperature on the detection and discrimination of tactile stimulation. Somatosens. Mot. Res. 14, 181–188. 10.1080/08990229771042.

21. Klinenberg, E., So, Y., and Rempel, D. (1996). Temperature effects on vibrotactile sensitivity threshold measurements: implications for carpal tunnel screening tests. J. Hand Surg. Am. 21, 132–137. 10.1016/S0363-5023(96)80166-3.

22. Thyagarajan, D., and Dyck, P.J. (1994). Influence of local tissue temperature on vibration detection threshold. J. Neurol. Sci. 126, 149–152. 10.1016/0022-510X(94)90265-8.

23. Ide, H., Akimura, H., and Obata, S. (1985). Effect of skin temperature on vibrotactile sensitivity. Med. Biol. Eng. Comput. 23, 306–310. 10.1007/BF02441583.

24. Weitz, J. (1941). Vibratory sensitivity as a function of skin temperature. J. Exp. Psychol. 28, 21–36. 10.1037/H0059426.

25. Harazin, B., Harazin-Lechowska, A., and Kałamarz, J. (2013). Effect of individual finger skin temperature on vibrotactile perception threshold. Int. J. Occup. Med. Environ. Health 26, 930–939. 10.2478/S13382-013-0163-6.

26. Frisina, R.D., and Gescheider, G.A. (1977). Comparison of child and adult vibrotactile thresholds as a function of frequency and duration. Percept. Psychophys. 22, 100–103. 10.3758/BF03206086/METRICS.

27. Verrillo, R.T. (1965). The effect of number of pulses on vibrotactile thresholds. Psychon. Sci. 37, 843–846. 10.3758/BF03343026.

28. Verrillo, R.T. (1965). Temporal Summation in Vibrotactile Sensitivity. J. Acoust. Soc. Am. 37, 843–846. 10.1121/1.1909458.

29. Gescheider, G.A., Hoffman, K.E., Harrison, M.A., Travis, M.L., and Bolanowski, S.J. (1994). The effects of masking on vibrotactile temporal summation in the detection of sinusoidal and noise signals. J. Acoust. Soc. Am. 95, 1006–1016. 10.1121/1.408464.

30. Green, B.G. (1977). The effect of skin temperature on vibrotactile sensitivity. Percept. Psychophys. 21, 243–248. 10.3758/BF03214234/METRICS.

31. Cheung, S.S. (2015). Responses of the hands and feet to cold exposure. Temperature: Multidisciplinary Biomedical Journal 2, 105. 10.1080/23328940.2015.1008890.

32. Isii, Y., Matsukawa, K., Tsuchimochi, H., and Nakamoto, T. (2007). Ice-water hand immersion causes a reflex decrease in skin temperature in the contralateral hand. J. Physiol. Sci. 57, 241–248. 10.2170/PHYSIOLSCI.RP007707.

33. Charkoudian, N. (2003). Skin blood flow in adult human thermoregulation: how it works, when it does not, and why. Mayo Clin. Proc. 78, 603–612. 10.4065/78.5.603.

34. Al, E., Iliopoulos, F., Forschack, N., Nierhaus, T., Grund, M., Motyka, P., Gaebler, M., Nikulin, V. V., and Villringer, A. (2020). Heart-brain interactions shape somatosensory perception and evoked potentials. Proc. Natl. Acad. Sci. U. S. A. 117, 10575–10584. 10.1073/PNAS.1915629117.

35. Grund, M., Al, E., Pabst, M., Dabbagh, A., Stephani, T., Nierhaus, T., Gaebler, M., and Villringer, A. (2022). Respiration, Heartbeat, and Conscious Tactile Perception. J. Neurosci. 42, 643–656. 10.1523/JNEUROSCI.0592-21.2021.

36. Galvez-Pol, A., Virdee, P., Villacampa, J., and Kilner, J.M. (2022). Active tactile discrimination is coupled with and modulated by the cardiac cycle. Elife 11. 10.7554/ELIFE.78126.

37. Saal, H.P., and Bensmaia, S.J. (2014). Touch is a team effort: interplay of submodalities in cutaneous sensibility. Trends Neurosci. 37, 689–697. 10.1016/J.TINS.2014.08.012.

38. Bell, J., Bolanowski, S., and Holmes, M.H. (1994). The structure and function of Pacinian corpuscles: a review. Prog. Neurobiol. 42, 79–128. 10.1016/0301-0082(94)90022-1.

39. Bolanowski, S.J., and Zwislocki, J.J. (1984). Intensity and frequency characteristics of pacinian corpuscles. I. Action potentials. J. Neurophysiol. 51, 793–811. 10.1152/JN.1984.51.4.793.

40. Mountcastle, V.B., LaMotte, R.H., and Carli, G. (1972). Detection thresholds for stimuli in humans and monkeys: comparison with threshold events in mechanoreceptive afferent nerve fibers innervating the monkey hand. J. Neurophysiol. 35, 122–136. 10.1152/JN.1972.35.1.122.

41. Mountcastle, V.B., Talbot, W.H., Dar-Smith, I., and Kornhuber, H.H. (1967). Neural basis of the sense of flutter-vibration. Science 155, 597–600. 10.1126/SCIENCE.155.3762.597.

42. Inman, D.R., and Peruzzi, P. (1961). The effects of temperature on the responses of Pacinian corpuscles. J. Physiol. 155, 280–301. 10.1113/JPHYSIOL.1961.SP006627.

43. Ishiko, N., and Loewenstein, W.R. (1961). Effects of temperature on the generator and action potentials of a sense organ. J. Gen. Physiol. 45, 105–124. 10.1085/JGP.45.1.105.

44. Tummala, N., Reardon, G., Dandu, B., Shao, Y., Saal, H.P., and Visell, Y. (2026). Biomechanical filtering supports efficient tactile encoding in the human hand. J. R. Soc. Interface 23. 10.1098/RSIF.2025.0793.

45. Gescheider, G.A., and Joelson, J.M. (1983). Vibrotactile temporal summation for threshold and suprathreshold levels of stimulation. Percept. Psychophys. 33, 156–162. 10.3758/BF03202833/METRICS.

46. Bochereau, S., Terekhov, A., and Hayward, V. (2014). Amplitude and duration interdependence in the perceived intensity of complex tactile signals. Lecture Notes in Computer Science (including subseries Lecture Notes in Artificial Intelligence and Lecture Notes in Bioinformatics) 8618, 93–100. 10.1007/978-3-662-44193-0_13/FIGURES/4.

47. Laeng, B., and Endestad, T. (2012). Bright illusions reduce the eye’s pupil. Proc. Natl. Acad. Sci. U. S. A. 109, 2162–2167. 10.1073/PNAS.1118298109.

48. Macmillan, N.A.., Hautus, M.J.., and Creelman, C. Douglas. (2022). Detection theory: a user’s guide (Routledge).

49. Okazawa, G., and Kiani, R. (2023). Neural Mechanisms That Make Perceptual Decisions Flexible. Annu. Rev. Physiol. 85, 191–215. 10.1146/ANNUREV-PHYSIOL-031722-024731.

50. Limbach, K., and Corballis, P.M. (2016). Prestimulus alpha power influences response criterion in a detection task. Psychophysiology 53, 1154–1164. 10.1111/PSYP.12666.

51. Zhou, Y.J., van Es, M.W.J., and Haegens, S. (2025). Distinct alpha networks modulate different aspects of perceptual decision-making. PLoS Biol. 23. 10.1371/JOURNAL.PBIO.3003461.

52. Carandini, M., and Heeger, D.J. (2011). Normalization as a canonical neural computation. Nat. Rev. Neurosci. 13, 51–62. 10.1038/NRN3136.

53. Rahman, M.S., and Yau, J.M. (2019). Somatosensory interactions reveal feature-dependent computations. J. Neurophysiol. 122, 5–21. 10.1152/JN.00168.2019.

54. Brouwer, G.J., Arnedo, V., Offen, S., Heeger, D.J., and Grant, A.C. (2015). Normalization in human somatosensory cortex. J. Neurophysiol. 114, 2588–2599. 10.1152/JN.00939.2014.

55. Vestergaard, M., Carta, M., Güney, G., and Poulet, J.F.A. (2023). The cellular coding of temperature in the mammalian cortex. Nature 2023 614:7949 *614*, 725–731. 10.1038/s41586-023-05705-5.

56. Schnepel, P., Paricio-Montesinos, R., Ezquerra-Romano, I., Haggard, P., and Poulet, J.F.A. (2024). Cortical cellular encoding of thermotactile integration. Curr. Biol. 34, 1718–1730.e3. 10.1016/J.CUB.2024.03.018.

57. Iggo, A., and Ramsey, R.L. (1976). Thermosensory mechanisms in the spinal cord of monkeys. Sensory Functions of the Skin in Primates, 285–304. 10.1016/B978-0-08-021208-1.50027-8.

58. Mariño, J., Martinez, L., and Canedo, A. (1999). Sensorimotor Integration at the Dorsal Column Nuclei. News Physiol. Sci. 14, 231–237. 10.1152/PHYSIOLOGYONLINE.1999.14.6.231.

59. Jones, E.G., and Powell, T.P.S. (1970). Connexions of the somatic sensory cortex of the rhesus monkey. 3. Thalamic connexions. Brain 93, 37–56. 10.1093/BRAIN/93.1.37.

60. Veldhuijzen, D.S., Greenspan, J.D., Kim, J.H., and Lenz, F.A. (2010). Altered pain and thermal sensation in subjects with isolated parietal and insular cortical lesions. Eur. J. Pain 14, 535.e1–535.e11. 10.1016/J.EJPAIN.2009.10.002.

61. Baier, B., Zu Eulenburg, P., Geber, C., Rohde, F., Rolke, R., Maihöfner, C., Birklein, F., and Dieterich, M. (2014). Insula and sensory insular cortex and somatosensory control in patients with insular stroke. Eur. J. Pain 18, 1385–1393. 10.1002/J.1532-2149.2014.501.X.

62. Birklein, F., Rolke, R., and Müller-Forell, W. (2005). Isolated insular infarction eliminates contralateral cold, cold pain, and pinprick perception. Neurology 65, 1381. 10.1212/01.WNL.0000181351.82772.B3.

63. Greenspan, J.D., Ohara, S., Franaszczuk, P., Veldhuijzen, D.S., and Lenz, F.A. (2008). Cold stimuli evoke potentials that can be recorded directly from parasylvian cortex in humans. J. Neurophysiol. 100, 2282–2286. 10.1152/JN.90564.2008.

64. Ostrowsky, K., Magnin, M., Ryvlin, P., Isnard, J., Guenot, M., and Mauguière, F. (2002). Representation of pain and somatic sensation in the human insula: a study of responses to direct electrical cortical stimulation. Cereb. Cortex 12, 376–385. 10.1093/CERCOR/12.4.376.

65. Craig, A.D., Chen, K., Bandy, D., and Reiman, E.M. (2000). Thermosensory activation of insular cortex. Nat. Neurosci. 3, 184–190. 10.1038/72131.

66. Ung, K., Magnotti, J.F., Kim, B., Yau, J.M., and Nordmark, P.F. (2025). Acute Loss of Tactile Input Leads to General Compensatory Changes in Eye-Hand Coordination during Object Manipulation. eNeuro 12. 10.1523/ENEURO.0487-23.2025.

67. Choi, C., Ma, Y., Li, X., Chatterjee, S., Sequeira, S., Friesen, R.F., Felts, J.R., and Cynthia Hipwell, M. (2022). Surface haptic rendering of virtual shapes through change in surface temperature. Sci. Robot. 7. 10.1126/SCIROBOTICS.ABL4543/SUPPL_FILE/SCIROBOTICS.ABL4543_SM.PDF.

68. Salvato, G., and Crucianelli, L. (2025). Shaping bodily self-awareness through thermosensory signals. Trends Cogn. Sci. 10.1016/J.TICS.2025.11.008.

69. Sakamoto, M., and Ifuku, H. (2021). Attenuation of sensory processing in the primary somatosensory cortex during rubber hand illusion. Sci. Rep. 11. 10.1038/S41598-021-86828-5.

70. Cordier, L., Fuchs, X., Herpertz, S., Trojan, J., and Diers, M. (2020). Synchronous Stimulation With Light and Heat Induces Body Ownership and Reduces Pain Perception. J. Pain 21, 700–707. 10.1016/J.JPAIN.2019.10.009.

71. Mosch, B., Fuchs, X., Tu, T., and Diers, M. (2025). Time course of the rubber hand illusion-induced analgesia. Pain Rep. 10. 10.1097/PR9.0000000000001252.

72. Benedetti, F., and Shaibani, A. (2025). A brief history of placebo. Handb. Clin. Neurol. 213, 3–15. 10.1016/B978-0-443-29884-4.00009-1.

73. Colloca, L. (2024). The Nocebo Effect. Annu. Rev. Pharmacol. Toxicol. 64, 171–190. 10.1146/ANNUREV-PHARMTOX-022723-112425.

74. Luo, Y., Lohrenz, T., Lumpkin, E.A., Montague, P.R., and Kishida, K.T. (2024). The expectations humans have of a pleasurable sensation asymmetrically shape neuronal responses and subjective experiences to hot sauce. PLoS Biol. 22, e3002818. 10.1371/JOURNAL.PBIO.3002818.

75. Trabanelli, S., Akselrod, M., Fellrath, J., Vanoni, G., Bertoni, T., Serino, S., Papadopoulou, G., Born, M., Girondini, M., Ercolano, G., et al. (2025). Neural anticipation of virtual infection triggers an immune response. Nat. Neurosci. 28, 1968–1977. 10.1038/S41593-025-02008-Y.

76. Oldfield, R.C. (1971). The assessment and analysis of handedness: the Edinburgh inventory. Neuropsychologia 9, 97–113. 10.1016/0028-3932(71)90067-4.

77. Jin, B.E., Wulff, H., Widdicombe, J.H., Zheng, J., Bers, D.M., and Puglisi, J.L. (2012). A simple device to illustrate the Einthoven triangle. Adv. Physiol. Educ. 36, 319–324. 10.1152/ADVAN.00029.2012.

78. Freeman, A.W., and Johnson, K.O. (1982). Cutaneous mechanoreceptors in macaque monkey: temporal discharge patterns evoked by vibration, and a receptor model. J. Physiol. 323, 21–41. 10.1113/JPHYSIOL.1982.SP014059.

79. Manfredi, L.R., Baker, A.T., Elias, D.O., Dammann, J.F., Zielinski, M.C., Polashock, V.S., and Bensmaia, S.J. (2012). The effect of surface wave propagation on neural responses to vibration in primate glabrous skin. PLoS One 7. 10.1371/JOURNAL.PONE.0031203.

80. Muniak, M.A., Ray, S., Hsiao, S.S., Dammann, J.F., and Bensmaia, S.J. (2007). The neural coding of stimulus intensity: linking the population response of mechanoreceptive afferents with psychophysical behavior. J. Neurosci. 27, 11687–11699. 10.1523/JNEUROSCI.1486-07.2007.

81. Johansson, R.S., and Vallbo, A.B. (1979). Tactile sensibility in the human hand: relative and absolute densities of four types of mechanoreceptive units in glabrous skin. J. Physiol. 286, 283–300. 10.1113/JPHYSIOL.1979.SP012619.

82. Torebjork, H.E., LaMotte, R.H., and Robinson, C.J. (1984). Peripheral neural correlates of magnitude of cutaneous pain and hyperalgesia: simultaneous recordings in humans of sensory judgments of pain and evoked responses in nociceptors with C-fibers. J. Neurophysiol. 51, 325–339. 10.1152/JN.1984.51.2.325.

83. Singh, G.S., and Acerbi, L. (2024). PyBADS: Fast and robust black-box optimization in Python. J. Open Source Softw. 9, 5694. 10.21105/JOSS.05694/STATUS.SVG).

84. Acerbi, L., and Ma, W.J. (2017). Practical Bayesian Optimization for Model Fitting with Bayesian Adaptive Direct Search. Adv. Neural Inf. Process. Syst. 2017*-December*, 1837–1847.

